# Motif Analysis in k-mer Networks: An Approach towards Understanding SARS-CoV-2 Geographical Shifts

**DOI:** 10.1101/2020.10.04.325662

**Authors:** Sourav Biswas, Suparna Saha, Sanghamitra Bandyopadhyay, Malay Bhattacharyya

## Abstract

With an increasing number of SARS-CoV-2 sequences available day by day, new genomic information is getting revealed to us. As SARS-CoV-2 sequences highlight wide changes across the samples, we aim to explore whether these changes reveal the geographical origin of the corresponding samples. The *k*-mer distributions, denoting normalized frequency counts of all possible combinations of nucleotide of size upto *k*, are often helpful to explore sequence level patterns. Given the SARS-CoV-2 sequences are highly imbalanced by its geographical origin (relatively with a higher number samples collected from the USA), we observe that with proper under-sampling *k*-mer distributions in the SARS-CoV-2 sequences predict its geographical origin with more than 90% accuracy. The experiments are performed on the samples collected from six countries with maximum number of sequences available till July 07, 2020. This comprises SARS-CoV-2 sequences from Australia, USA, China, India, Greece and France. Moreover, we demonstrate that the changes of genomic sequences characterize the continents as a whole. We also highlight that the network motifs present in the sequence similarity networks have a significant difference across the said countries. This, as a whole, is capable of predicting the geographical shift of SARS-CoV-2.

## 1 Introduction

We have already seen how the COVID-19 pandemic, originating from Wuhan in China [1], has spread to more than 200 countries and territories around the world. This disease has already been characterized by the identification of a novel type of betacoronavirus, termed as SARS-CoV-2, which primarily infects human [2]. There have been various strains of this virus that are prevalent across the world [3, 4]. However, it is still unclear how the changes in SARS-Cov-2 sequences characterize the spread of different strains in multiple countries.

Coronaviruses are a large family of RNA viruses that cause diseases in mammals and birds [5]. Coronaviruses have been identified broadly in avian hosts along with various mammals, which include camels, masked palm civets, bats, mice, dogs, cats, and humans. The coronaviruses cause respiratory and neurological diseases in hosts [6]. In humans, it causes several illnesses, ranging from the mild common cold to many severe diseases such as SARS-CoV (Severe Acute Respiratory Syndrome) and MERS-CoV (Middle East Respiratory Syndrome). They are enveloped viruses with a positive-sense single-stranded RNA genome. The genome size of coronaviruses ranges between 26-32 kb [7], the largest among known RNA viruses [8].

The sequencing of viral genomes has opened up new promises through bioinformatics analyses [9, 10]. With the recent efforts of sequencing the COVID-19 samples collected across the world, thousands of genomic samples are becoming available day by day [11]. This has lead to the discovery of new pathophysiological facts about coronaviruses. It is equally interesting to explore the number of mutations happening in the samples emerging across the world. In this paper, we aim to verify whether the changes of sequence in the samples can reveal the geographic location of this virus. We basically adopt a network motif analysis approach for the said purpose.

## 2 Related Work

In earlier studies, it was strongly suggested that the origin of SARS-CoV was SARSr-CoV in bats [12, 13, 14]. In fact, their sequence similarity highlights discordant clustering with the Bat-SARS-like coronavirus sequences [15]. The S and ORF8 genes are the most varying genomic regions in SARS-CoV and SARSr-CoV [16, 17], where the S gene is important for receptor binding in SARS-CoV infection and is highly alterable in the deletions of some amino acids in the S protein [13, 18, 19, 20]. Based on the changes of amino acids, 3 categories of Coronaviruses have been characterized, termed as A, B, and C. A is the ancestral category of bat outgroup coronavirus. Type A and C are significantly found outside of East Asia whereas the type B is prevalent in East Asia. The phylogenetic network helps to trace the route of COVID-19, can prevent the worldwide spread of the disease.

The spike (S) protein performs a very influential role in the attachment and entry of the virus, and is responsible for the development of antibodies and vaccines. Tai et al. identified the receptor-binding domain (RBD) in SARS-CoV-2 S protein [21]. Moreover, they found that RBD protein binds to human and bat angiotensin-converting enzyme 2 (ACE2) receptors intensely. Notably, RBD protein showed a remarkably higher association with ACE2 than SARS-CoV RBD. Tai et al. indicated that specific antibodies may be used for the treatment of SARS-CoV-2 infection and characterized SARS-CoV RBD as a candidate vaccine to produce cross-reactive antibodies for the prevention of SARS-CoV-2. Though the RBD of SARS-CoV-2 differs widely from the SARS-CoV at the C-terminus, such a difference will not make drastic changes while associating with the ACE2 receptor but had an impact on the cross-reactivity of the antibodies.

Another well-defined approach by Dong et al. measures the distance between both genomic and proteomic data, and suggested that pangolins are the most probably intermediate host of the SARS-CoV-2 outbreak[22]. There are significant variations in the bat coronavirus than in pangolin-cov compared with SARS-CoV-2 in the alignments of the spike surface glycoprotein receptor binding domain. Zhang et al. identified the pangolin as a missing link in the transmission of SARS-CoV-2 from bats to humans [23]. One of the recent analyses also revealed that SARS-CoV-2 expresses itself an isolated homogeneous cluster with one of the closest isolate followed by a group of Pangolin-CoVs [24].

Human-to-human transmission among very close contacts can spread out coronavirus gradually in the community. Li et al. performed a fundamental assessment of the transmission dynamics and epidemiologic characteristics of Pneumonia infected by COVID-19 [25]. It was found through appropriate statistical analysis that the basic reproductive number (*R*_0_) of COVID-19 is 2.2. In a very recent study by Yang et al., four genetic clusters of corona virus were identified [26]. These are mainly responsible for the major outbreaks defined as super spreaders.

There is an important impact of the age structure in the population for the fatality rates. As deaths are mostly seen at older ages, a recent report highlighted the vital role of demography [27]. The age structure helps to explain the death rates varies across the countries as well as unfolds the transmission mechanism. However, as of now, there is no antiviral treatment or vaccine. Hence, there is a serious need for safe as well as effective therapeutics against SARS-CoV-2 infection to save our world population.

There have been several attempts by the researchers to understand the footprint of the SARS-CoV-2 virus. Most of these analyses are either sequence-based or phylogenetic tree-based. Biswas et al. first showed that *k*-mer distributions in the earlier SARS-Cov-2 sequences can predict its collection timeline with a reasonable accuracy [28]. However, such patterns disappear over time. This may be because of the reason that the virus has spread over multiple countries and has acquired region-specific mutations. Forster et al. performed a phylogenetic network analysis with 160 complete genomes of SARS-CoV2 or SARS-CoV-2 [29].

However, there is hardly any effort to characterize the sequences based on their location of origin. In this paper, we show how we can predict the location of a SARS-CoV-2 sample from its sequence patterns and network motifs.

## 3 Materials and Methods

The materials and methods used in this paper are all available either from the experimenters through public repositories or from us for public use. Additional details are given below.

### 3.1 Dataset Details

The SARS-CoV-2 genomic sequences were collected from the NCBI Virus portal^1^ that cover multiple countries and continents. There are over eight thousand genome sequences on the host as humans, including complete and some incomplete genome. To date, only a few complete genome data are available for SARS-CoV-2 from hosts other than humans for example, (Canis lupus familiaris^2^). We limited our findings only to human as a host in this study. A complete sequence provides the whole genome data which has a basepair length around 30 kilobases. In order to get the most information out of it, we took those gene sequences which are complete. This approach narrows down to 6262 sequences excluding the RefSeq sequence (Accession no NC_045512) which are attached as a supplementary file **6262 sequences_7thjuly** (dataset 1). A few of these sequences have missing collection dates, hence we pre-processed the raw data by omitting those missing values. Now the number of samples becomes 5817 which is included in the supplementary file **kmer_result_5817_kmer_values**(dataset 2). The Table 1 shows the samples (total 5817) for all countries (total 45 countries) which is highly imbalanced. The USA has the highest number of samples of 4323, whereas the second highest Australia has 444 samples; and many of them are even below 20. Among these 45 countries, the top six classes (Australia, China, India, Greece, France and USA) were chosen for countrywise classification. These six countries were selected based on a cut-off (having at least eighty samples) that retains a good geographical diversity. The samples from all the countries are used for continentwise classification.

**Table 1:** Countrywise distribution of total 5817 SARS-CoV-2 nucleotide sequences available in NCBI as on July 07, 2020.

| Country | Percentage | # Samples | Time Gap between Sample Collection and Sequence Release (Weeks) |
| --- | --- | --- | --- |
| **USA** | 74.3166581 | 4323 | 5.752 |
| **Australia** | 7.63280041 | 444 | 6.637 |
| **India** | 5.44954444 | 317 | 2.578 |
| **China** | 1.7019082 | 99 | 10.009 |
| **Greece** | 1.66752622 | 97 | 8.080 |
| **France** | 1.39247035 | 81 | NA |
| **Bangladesh** | 1.30651539 | 76 | 2.943 |
| **Egypt** | 1.2721334 | 74 | NA |
| **Germany** | 0.79078563 | 46 | 12.868 |
| **Taiwan** | 0.53292075 | 31 | 7.392 |
| **Spain** | 0.41258381 | 24 | 3.366 |
| **Czech Republic** | 0.39539281 | 23 | 7.385 |
| **Rest of the 33 Countries** | 3.1287 | 182 | NA |
| **Total** | **100** | **5817** |  |

### 3.2 Methods

Since the dataset is highly imbalanced (USA alone holds 74% of the total sequences), we preprocessed the data before classification. Note that, different countries publish their sequence data with different time gaps from the date of sample collection. The fourth column of Table 1 highlights the mean time difference (in weeks) between the collection of samples and release of the same. However, it is not a comparable measure across the countries since many of the countries have released limited number of samples so far. However, this strongly suggests that it is principally better to select the collection months (instead of release months) for the prediction of its origin (country or continent). Based on this understanding, we segregated the samples according to their collection months (considering a bin size of 30 days).

For each month of collection, we took the top countries that had more genomic data (sequences of SARS-CoV-2 samples) than the others. Before applying any classification approach, data was turned into a balanced form. For this, we performed random stratified under-sampling on the data (samples from corresponding countries) suffering from imbalance to a count same as the average of other countries. This is separately done for each of the classification tasks associating all the data. More details are given in the Results section. Note that, the basic models like linear regression is not suitable for a multi-class classification problems, hence we did not try them out. We performed the continent classification using the data collected over all samples irrespective of their collection months.

For each of the above classification results, we have performed a comprehensive analysis using the network motifs. A small connected induced subgraph of a network is called a motif if it appears significantly higher in count in that network in comparison with similar random networks [30]. If the frequency of a subgraph is lower than that of the similar random networks it is defined as an anti-motif [31]. Both are important for the analysis of topological patterns of a network. We used the network motif detection tool FANMOD [32] for identifying significant substructures in the sequence similarity network. A sequence similarity network, in the context of the current paper, symbolizes the SARS-CoV-2 samples as nodes and a significant similarity (measured in terms of small distance) between them as an edge. Fig. 1 shows the conventional motif or graphlet ID generation method from corresponding adjacency matrix. It is simply the decimal number when converted its adjacency binary matrix to a binary string following row-major order.

**Figure 1:**
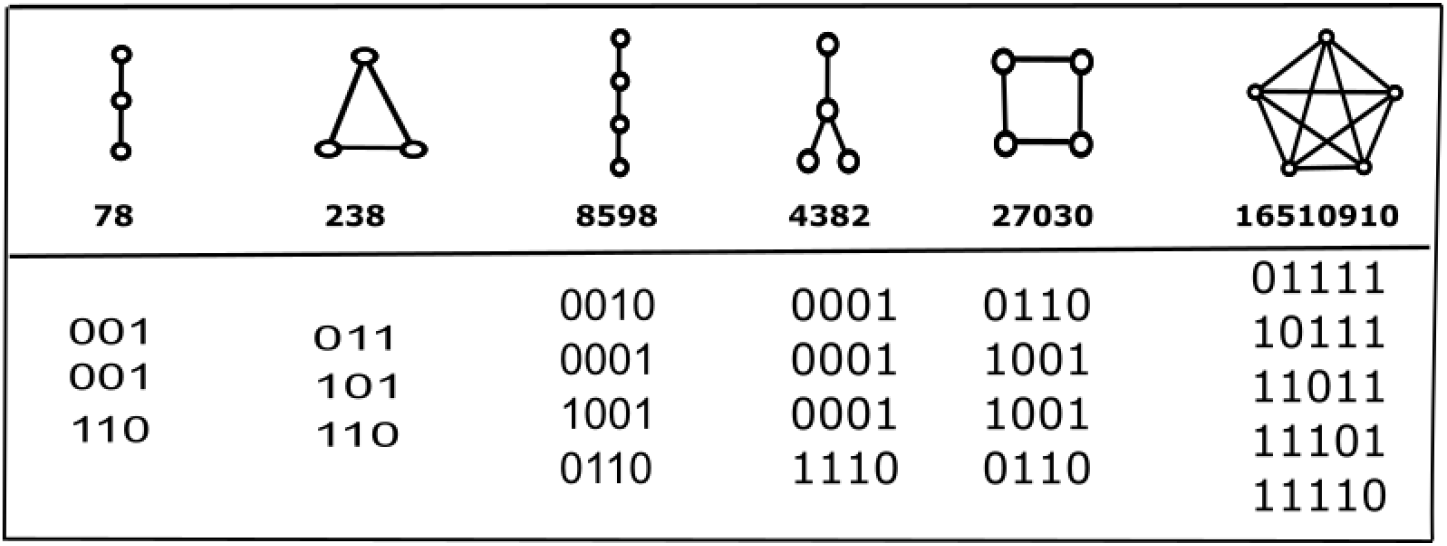
Example of sample motif orgraphlet ID generation from the corresponding adjacency matrix.

The motivation behind the motif analysis is that it is not effected by the imbalance class datasets. To build the *k*-mer-based network from the *k*-mer coverage, we first calculated a distance matrix (using package ‘philentropy’ in R, method = ‘Euclidean Distance’) and made it a binary adjacency matrix by taking the median as a cutoff point. The distance function finds the distances between pairwise *k*-mer distribution of each genome sequence sample. We include an edge between a pair of nodes (corresponding to SARS-CoV-2 sample sequences) their distance fall under a defined cutoff value. This finally yields a binary adjacency matrix. By following the same approach, we created *k*-mer-based network for each countries as it was in the *k*-mer coverage classification. In the result section we reported the motifs that we found with sizes 3, 4 and 5, which supports our hypothesis of geographical signatures of SARS-CoV-2 sequences. It is shown that the networks build from the *k*-mer distributions of different countries are separable.

Several classification algorithms were performed that are standard [33]. The classification results and visualizations were obtained using *Orange*^3^ and R 4.0.1 in Windows^4^. The values of the parameters of different algorithms are set as required to overcome the underfitting and/or overfitting of the models in *Orange*. The performance of the classification models are determined in terms of the five evaluation criteria, namely 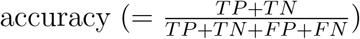, 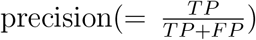, 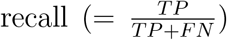, F1 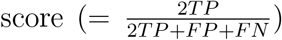, and AUC (area under ROC curve), where *TP, TN, FP* and *FN* stands for the number of true positives, true negatives, false positives and false negatives, respectively.

## 4 Results

We conducted analytical experiments on the samples collected until July 07, 2020 from several countries. The frequency counts for several *k*-mers (for *n* =1, 2 and 3) were extracted yielding a total 84 (4+ 16 + 64) features. The frequency values were normalized (by dividing with the length of the sequence). Since considering higher *k* value yield exponentially high computational cost and a codon consists only three nucleotides, we choose *k* to be limited up to 3.

### 4.1 *k*-mer Analysis and Classification Results

We first demonstrate how the coverage of *k*-mers can help to identify the geographical origin of a SARS-CoV-2 sample. It is found that with enough number of samples available from each geographical locations classification results are very good. We first tried this on a few countries and then on the continents. *K*-mer distribution produces good accuracy with different classifiers in the both cases, which are explained in the following two sub-sections.

#### 4.1.1 Classification of Countries

We choose the top six countries as they all have at least 80 and more samples and are from different diverse parts of the world. Hence these countries as Australia, China, India, Greece, France, and USA belongs to major continents with vast geographical area. As USA has the most unbalanced number of samples we used the random stratified under-sampling method. The number of USA samples have now reduced to 260, which is the average number of samples of the rest five countries. The figure 2 shows the frequency distributions of the samples across these six countries after under-sampling. We applied four classifiers as kNN, RandomForest, NaiveBayes, and Constant in two different approaches: 10-fold cross validation and 70-30% splitting of dataset (Train and Test). The Table 2 and the table 3 shows the result of these two approaches. In both the cases, RandomForest stands out to get the best outcomes in terms of all performance measures of different classifiers. We also used a *Costant classifier* (which means no classifier in reality) only to compare it with other classifiers. We saw that the accuracy falls with this approach thereby indicating that the real classifiers are not over-fitting. Therefore, the results are indeed significant based on geographical diversity.

**Figure 2:**
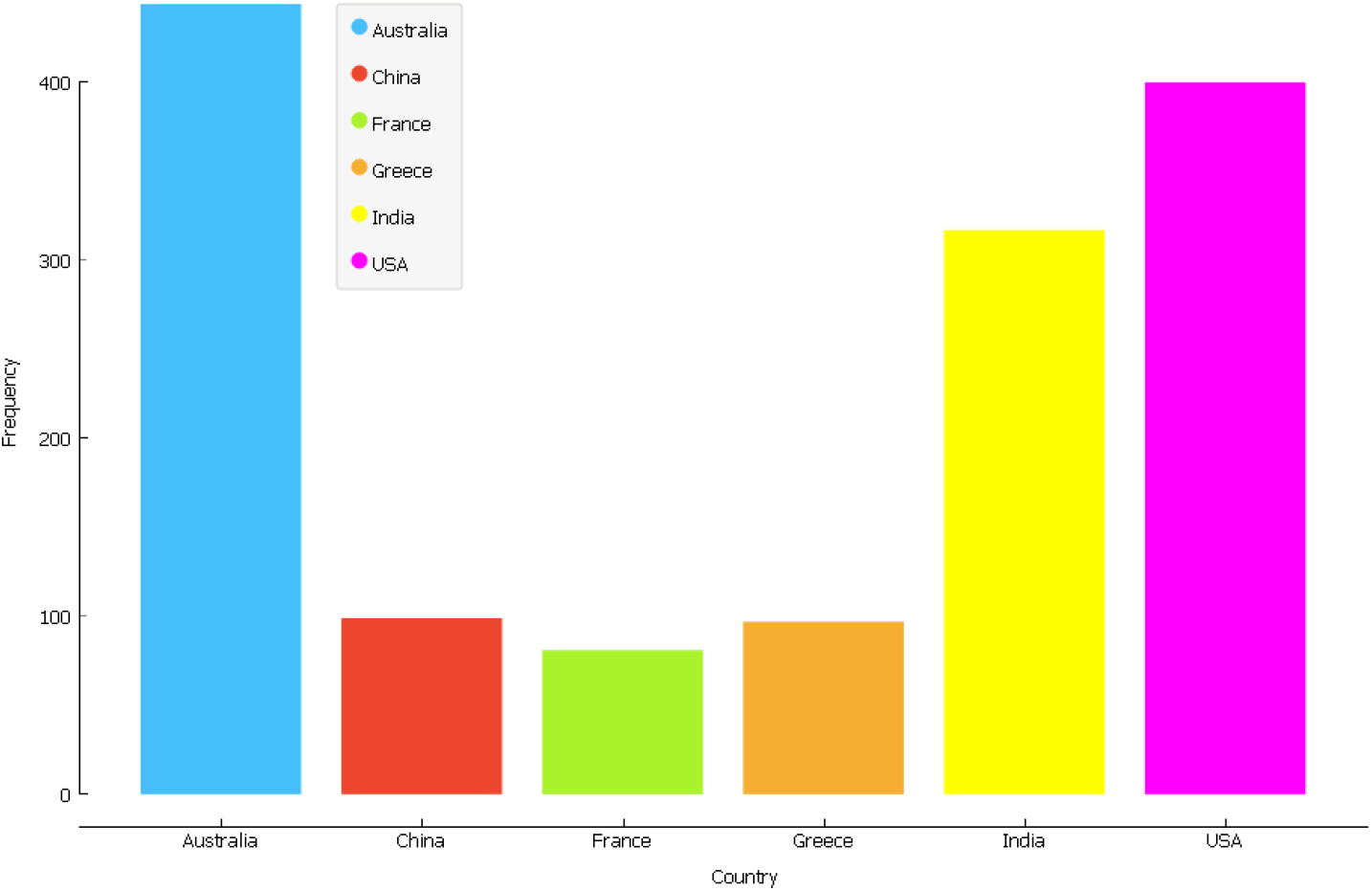
Distribution of samples across the six countries Australia, China, France, Greece, USA, and India.

**Table 2:** Performance evaluation of different classification approaches (stratified 10-fold cross validation) based on the 84 *k*-mer features to distinguish the countries from where samples were collected. The results are approximated upto 3 decimal places. The best values over a column are shown in bold.

| Model | AUC | CA | F1 Score | Precision | Recall |
| --- | --- | --- | --- | --- | --- |
| kNN | 0.964 | 0.869 | 0.868 | 0.872 | 0.869 |
| Random Forest | **0.994** | **0.937** | **0.937** | **0.938** | **0.937** |
| Naive Bayes | 0.968 | 0.831 | 0.835 | 0.855 | 0.831 |
| Constant | 0.495 | 0.342 | 0.174 | 0.117 | 0.342 |

**Table 3:** Performance evaluation of different classification approaches with separate train and test data based on the 84 *k*-mer features to distinguish the countries. Training and test data are taken as 70% and 30%, respectively.

| Model | AUC | CA | F1 Score | Precision | Recall |
| --- | --- | --- | --- | --- | --- |
| kNN | 0.953 | 0.838 | 0.838 | 0.853 | 0.838 |
| Random Forest | 0.991 | 0.931 | 0.930 | 0.932 | 0.931 |
| Naive Bayes | 0.965 | 0.830 | 0.837 | 0.863 | 0.830 |
| Constant | 0.5 | 0.339 | 0.172 | 0.115 | 0.339 |

#### 4.1.2 Classification of Continents

We grouped the different samples into the corresponding continents. This is done for all the countries irrespective of the count if samples they represent. The distribution of samples across different continents are shown in Table 4. ‘Out of the total 5 continents (Asia, Africa, Europe, America and Oceania), we exclude Africa for too low number of sequences and consider the rest continents. Again, here Americas has huge samples which makes the dataset unbalanced. Hence, we did random under-sampling of the continent and choose 460 samples (which is the average of the other three continents). After this pre-processing we have total 1839 samples (Americas 460 and the rest as per the table 4, excluding 84 sequences from Africa) which is used for the classification. This raw data is added as supplementary file **1839_data_USA_undersampled_excluding_Africa**(dataset 3).

**Table 4:** Continentwise distribution of total 5817 SARS-CoV-2 nucleotide sequences after pre-processing.

| Continent | # Samples |
| --- | --- |
| Africa | 84 |
| Americas | 4354 |
| Asia | 612 |
| Europe | 319 |
| Oceania | 448 |
| Total | 5817 |

The Table 5 shows the result of classification of the samples with 10-fold cross validation. For comparison we keep the constant classifier, which randomly assigns samples to classes. The poor performance of constant proves that the models are not over-fitting.

**Table 5:** Classification result on continent data with 10-fold cross validation.

| Model | AUC | CA | F1 Score | Precision | Recall |
| --- | --- | --- | --- | --- | --- |
| kNN | 0.945 | 0.825 | 0.826 | 0.828 | 0.825 |
| Random Forest | 0.983 | 0.903 | 0.902 | 0.903 | 0.903 |
| Naive Bayes | 0.916 | 0.717 | 0.714 | 0.750 | 0.717 |
| Constant | 0.498 | 0.333 | 0.166 | 0.111 | 0.333 |

#### 4.1.3 Classification of selected countries in March and April

As the genomic sequence data of SARS-CoV-2 has gradually increased over the last few months, we divided the collected samples into six different bins based on their month of collection, ranging between January 2020 and June 2020. Fig. 3 shows the proportion of distribution of genomic data available for the top six countries over the period of January to June, 2020. It can be seen from this figure that most of the sequences come from March and April, 2020. except for India and China. The reason behind is that, it started in China first and in India the infection started to peak later. Hence, we took the sequences from the four countries (Australia, France, Greece and USA) and the three countries (Australia, India and USA) in March and April respectively. The 10-fold cross validation results are shown in Table. 6 and Table. 7, respectively. As can be seen from these tables, classification accuracy values are quite high when the count of samples are well distributed among the countries of origin.

**Figure 3:**
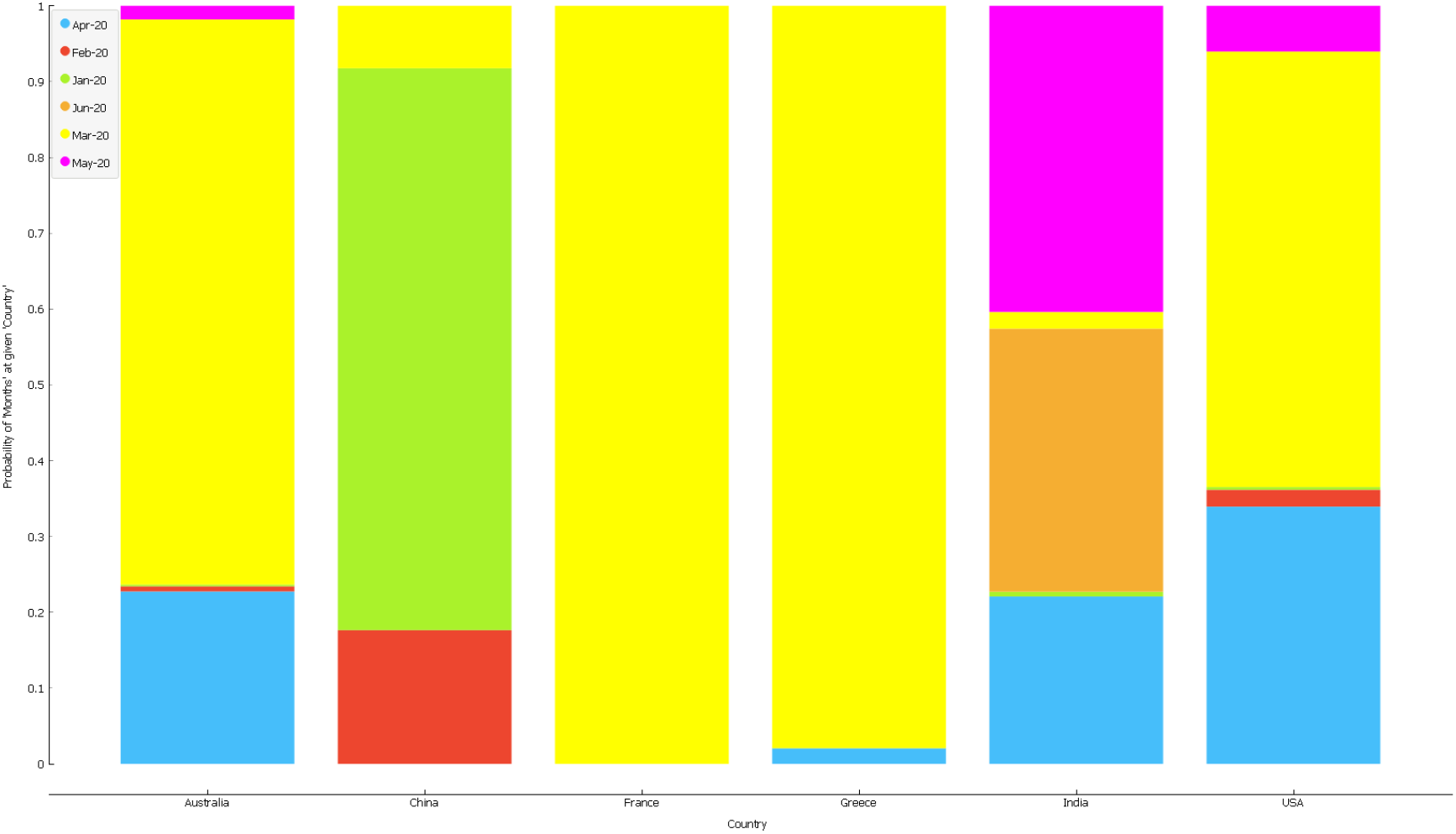
Monthwise distribution of samples collected from the top six countries.

**Table 6:** Classification results with 10-fold cross validation of the countries (Australia, USA, Greece and France) based on the SARS-CoV-2 samples collected from there in March 2020.

| Model | AUC | CA | F1 Score | Precision | Recall |
| --- | --- | --- | --- | --- | --- |
| kNN | 0.973 | 0.899 | 0.897 | 0.901 | 0.899 |
| Random Forest | 0.991 | 0.945 | 0.945 | 0.946 | 0.945 |
| Naive Bayes | 0.977 | 0.885 | 0.887 | 0.897 | 0.885 |
| Constant | 0.493 | 0.490 | 0.322 | 0.240 | 0.490 |

**Table 7:** Classification results with 10-fold cross validation of the countries (Australia, India and USA) based on the SARS-CoV-2 samples collected from there in April 2020.

| Model | AUC | CA | F1 Score | Precision | Recall |
| --- | --- | --- | --- | --- | --- |
| kNN | 0.930 | 0.844 | 0.844 | 0.848 | 0.844 |
| Random Forest | 0.987 | 0.934 | 0.934 | 0.936 | 0.934 |
| Naive Bayes | 0.954 | 0.852 | 0.85 | 0.861 | 0.852 |
| Constant | 0.488 | 0.393 | 0.222 | 0.154 | 0.393 |

### 4.2 Network Motif Analysis

As described earlier in the method section, network motifs can be thought of the building blocks of a network. This is a topological feature of a network but it reflects functionality also. Here, we have the same dataset used as in classifications of countries using *k*-mer distribution. Next, we build countrywise network based on the *k*-mer distribution. We built a pairwise distance matrix (Eucledian distance) of a country’s sequences. If it has *n* number of samples, the output would be a *n* * *n* square matrix. To reduce the high computational usage of motif detection, we limited our study to random 100 samples or less (since France and Greece have less than 100) from each country. We took the median for each of distance matrix and transform it into a binary matrix by using it as a cutoff value. Distance values less than cutoff are replaced by one (TRUE) and the rest are zeros (FALSE). Now the binary distance matrix can be regarded as the adjacency matrix of the corresponding country’s network. We hereafter refer to this as *k*-mer network. We applied the FANMOD tool [32] to explore the motifs. Among the possible 29 isomorphic graphlets/subgraphs (see Fig. 4), we found a range of motifs from 8 to 17 in the *k*-mer networks of those six countries. Most of the countries have around 8 motifs with only Greece as an exception having 17 motifs (see Table 8). The corresponding motif structure can be found in Fig. 4. f=For example, index *k* in Table 8 represents the topological structure of the corresponding motif as G_k_ in Fig. 4. Note that, there is no ordering of 29 graphlets shown in Fig. 4, hence it is not unique or universal like the motif IDs. The number of overlapping motifs among the six networks is eight (motif IDs as reported in the Table 8. The structures associated with the motif-IDs are also given as G1, G4, G5, G14, G19, G21, G25 and G27 according to the Fig. 4.

**Figure 4:**
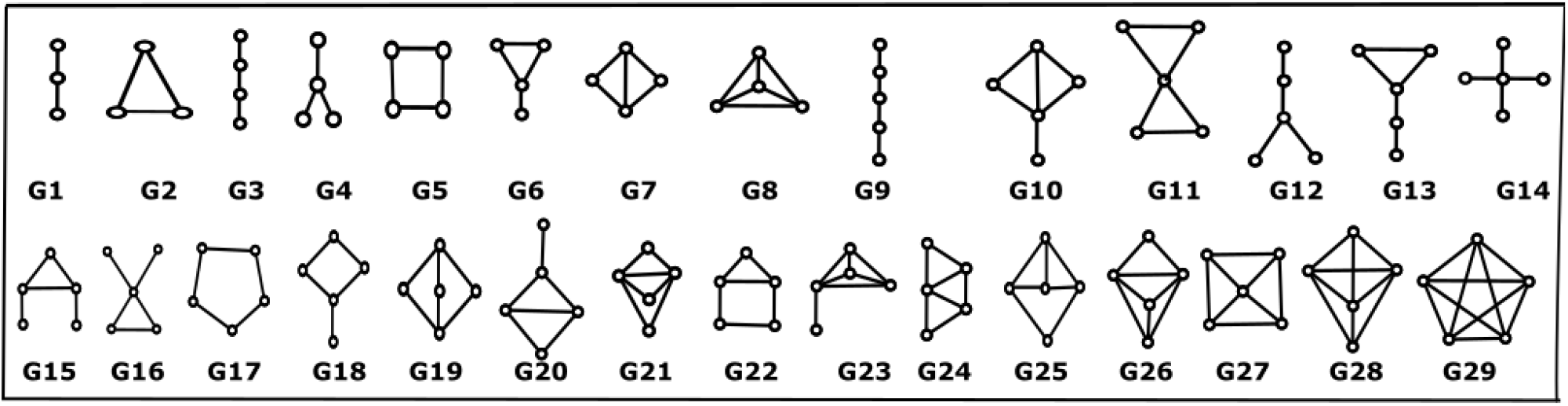
All possible 29 undirected graphlets of size 3, 4 and 5.

**Table 8:**
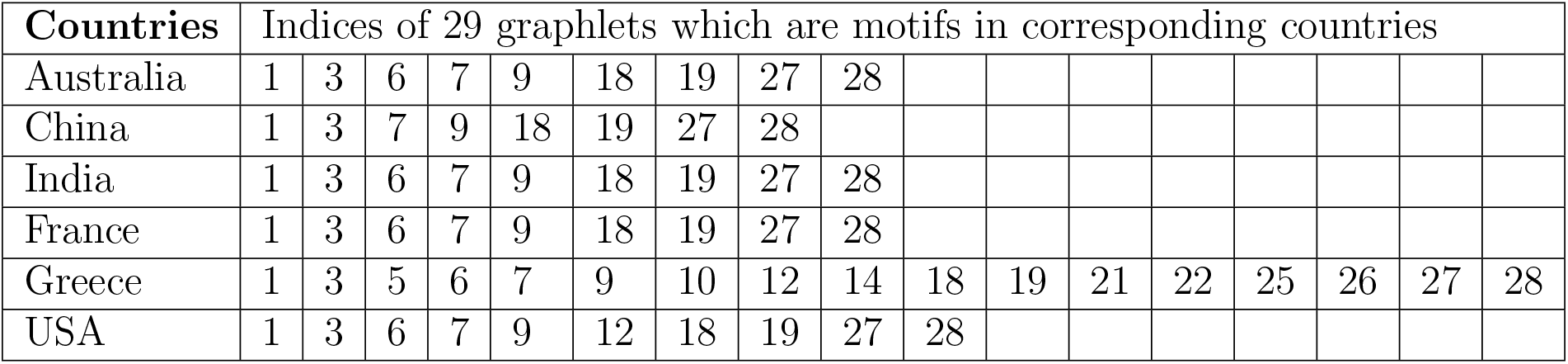
Distibution of network motifs across different countries. While some of the motifs are common across all the countries, several others are prsent in limited countries showing a distinctive nature.

**Table 9:** Z-scores of motifs of size 3, 4 and 5 found in the *k*-mer networks having a p-value < 0.05. The corresponding motif IDs and motif indices are reported.

| Countries | G1 | G4 | G5 | G14 | G19 | G21 | G25 | G27 |
| --- | --- | --- | --- | --- | --- | --- | --- | --- |
| Australia | 92.425 | 55.41 | 202.06 | 50.402 | 617.03 | 24.098 | 63.242 | 291.94 |
| China | 17.126 | 17.063 | 19.571 | 21.367 | 34.29 | 8.113 | 13.109 | 33.938 |
| India | 3.192 | 3.2924 | 3.7423 | 3.8945 | 4.7831 | 2.3526 | 5.6385 | 4.8308 |
| France | 27.087 | 30.234 | 48.21 | 32.352 | 121.85 | 14.182 | 27.018 | 61.019 |
| Greece | 20.733 | 29.694 | 57.31 | 31.355 | 225.43 | 24.081 | 59.535 | 60.626 |
| USA | 35.796 | 39.616 | 73.031 | 43.04 | 260.3 | 22.811 | 95.152 | 81.734 |

#### 4.2.1 Geographical *k*-mer network analysis by network motifs

There are six *k*-mer networks of six countries representing sequence similarities among them based on *k*-mer distribution. Each node of the network stands for a virus genome and the edges represent the similarity between two nodes. Similarity values are binary, either similarity exists or does not. The motifs were found (using FANMOD, [32]) with comparing the six countries’ networks with similar hundred randomized networks (generated for each country by FANMOD). The motifs or ‘the topological representation of sequential similarity’ of a particular country’s genome are not random as it would be for similar a hundred random networks.

Each motif represents the sequential similarity (*k*-mer distribution) within the genome samples for each of the countries. All motifs (except G1 and G5) have a common topological tendency such as, hub nodes with high degree/high betweenness centrality. Except G5, no ring structure are found which explains virus samples go for random mutation and the do not converge to a previous one by any way. The divergence of the genomes were found in all six countries. Notably, the graphlets that are found as motifs have resembling topology.

We further consider the z-scores of the motifs as a feature vector of a country and did a hierarchical clustering. The dendrogram and the heatmap are shown in Fig. 6 and 5. Since data were taken over the months for each country, it extracts the geographical shifts of the SARS-CoV-2. Two Asian countries India and China are grouped together. France emerges as in between two sets. USA and Greece have similarity for unknown reason where Australia has a completely different class in the dendrogram, which supports the diversity of genome samples over geographical shifts. The heatmap also justifies the fact of diversity of viral samples across the countries.

**Figure 5:**
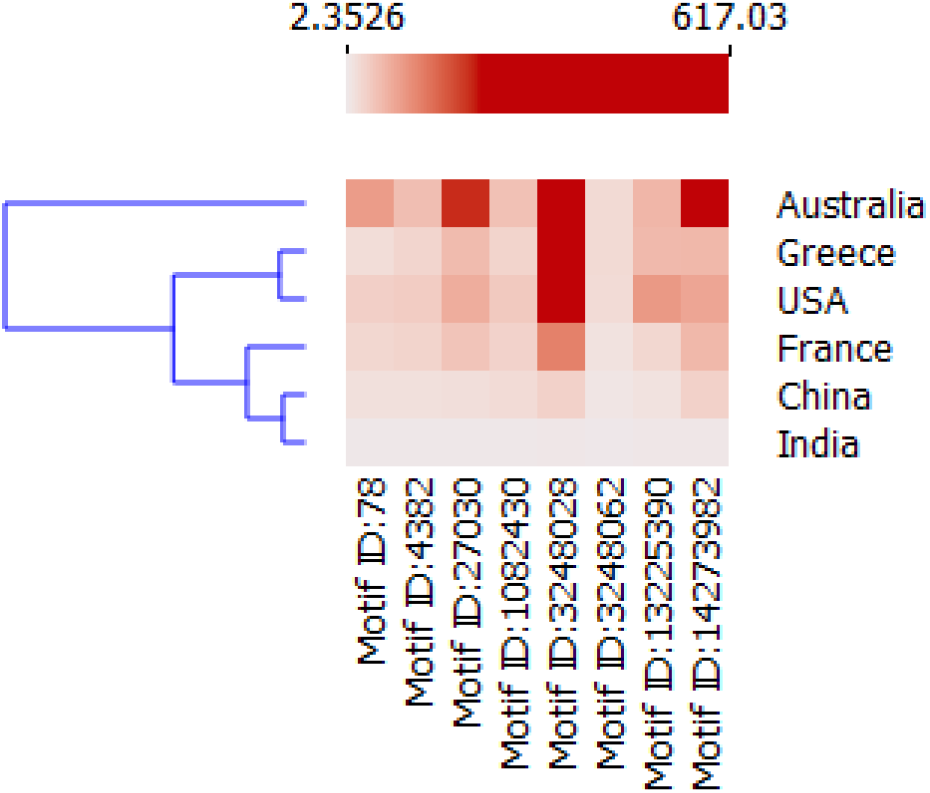
Heatmap of the z-scores obtained for the six countries.

**Figure 6:**
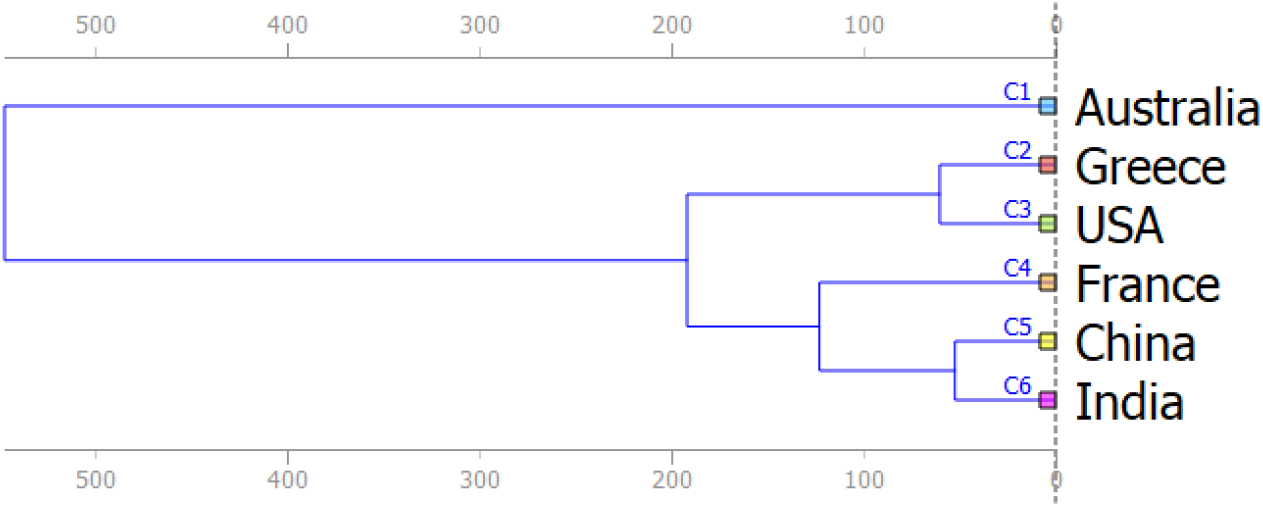
Hierarchical clustering with weighted linkage method on Euclidean distance matrices to group the countries based on the SARS-CoV-2 samples collected.

## 5 Conclusion

Our results demonstrate that the sequence alterations in SARS-CoV-2 samples collected across diverse geographical location provides distinctive patterns to predict its origin. These alterations are principally indebted to the environmental factors. We showed that with appropriate under-sampling, we can distinguish the countries and even the continents given the SARS-Cov-2 sequences of the samples collected from there. This highlights the existence of unique strains across different geographical regions. We also attempted to classify the samples by some countries across the period (March and April, 2020) such that the sequences are well distributed. We observed a high distinctive pattern of the sequences without adopting any sampling strategy. This indeed ensures the importance of the patterns explored through *k*-mer distribution. However, it is important to note that, though the sequences can be put within monthly bins collected over different timelines, different countries publish their data with different time gaps (after sample collection). Hence, this kind of classification task is really challenging. Biologically speaking, our results indicate that the mutations happening in SARS-CoV-2 over the time and place hold important clues to understand its evolution. Interestingly, as the countries and continents are distinguishable from the changes in SARS-CoV-2 sequences and network motifs, there lies a promised to adopt a genomic fingerprinting approach to better understand the footprint of this virus.

## Abbreviations

SARS-CoV-2: 2019 novel coronavirus
SARS-CoV: Severe Acute Respiratory Syndrome
MERS-Cov: Middle East Respiratory Syndrome
COVID-19: 2019 novel coronavirus disease

## Consent for publication

Not applicable.

## Ethics approval and consent to participate

Not applicable.

## Availability of data and materials

All datasets used in this paper are freely available from the NCBI data repository. The metadata for all the 6262 sequences (collected on 7th July from NCBI) is provided as Supplementary Dataset 1. The dataset containing 5817 sequences (after preprocessing) with normalized *k*-mer values is provided as Supplementary Dataset 2. The dataset containing 1839 samples (after undersampling the samples from USA and excluding Africa), with *k*-mer values used for continent classification, is provided as Supplementary Dataset 3.

## Authors’ contributions

SoB, SS, and MB designed the experiments. SoB and SS carried out the experiments and data analysis. SoB, SS, SaB, and MB drafted the manuscript. All the authors read and approved the final manuscript.

## Acknowledgments

SoB acknowledges Digital India Corporation (formerly Media Lab Asia), Ministry Of Electronics and Information Technology (MeitY), Government of India, for providing him a Senior Research Fellowship under the Visvesvaraya Ph.D. Scheme for Electronics and IT.

## Condolence

The authors feel deep grief and sympathy for those who got affected by the SARS-CoV-2

## Footnotes

1 https://www.ncbi.nlm.nih.gov/labs/virus (accessed on July 7th, 2020)

2 https://www.ncbi.nlm.nih.gov/nuccore/MT270814

3 https://orange.biolab.si

4 https://cran.rstudio.com/bin/windows

